# Phenotypic screen-based discovery of a small molecule that can increase adult neurogenesis and improve memory

**DOI:** 10.1101/2023.07.27.550845

**Authors:** Julie Davies, Anna Hoerder-Suabedissen, Ksenia Musaelyan, Blanca Torroba, Jesse Daubney, Nicole Untermoser, Tom Carter, Ulrich Bauer, Roderick Walker, Kate S. Harris, Liam Bromhead, Megalakshi Suresh, Penelope Fouka, Yichen Li, Steve Davies, Caleb Webber, David Bannerman, Georg Terstappen, Angela Russell, Francis G. Szele

## Abstract

Stem cells and neurogenesis persist in the postnatal and adult brain. Adult brain stem cells can be neuroprotective in disease and augment hippocampal-dependent cognitive function and thus are an important therapeutic target. Although many molecules have been discovered that regulate neurogenesis, few studies have attempted to amplify the process pharmacologically as a therapeutic goal. To address this gap, we used murine neurosphere cultures from the two major stem cell niches: the subventricular zone (SVZ) and the subgranular zone (SGZ). We screened compounds sharing pharmacophores with known inducers of neurogenesis and found several dozen proneurogenic compounds in an *in vitro* phenotypic screen. One, OXS-N1 was stable, and had acceptable absorption, distribution, metabolism, and excretion profiles in animal studies. OXS-N1 could increase neurogenesis in the SVZ and SGZ in WT mice after both intraperitoneal and oral administration. The number of newborn neurons (BrdU+/NeuN+) was increased; however, the number of activated stem cells (BrdU+/GFAP+) was not, suggesting an effect on neurogenesis independent of stem cell activation. This was supported by OXS-N1 increasing neurosphere differentiation but not proliferation. OXS-N1 also increased neurogenesis and improved performance in a Y maze cognitive task in PDGF-APPSw,Ind mice, a model of Alzheimer’s disease. RNAseq of SVZ and SGZ neurospheres in turn showed that genes associated with synaptic function were significantly increased by OXS-N1. Our study demonstrates the utility of phenotypic screening for the identification of molecules that increase neurogenesis and might be of therapeutic relevance.

## Introduction

Neurogenesis is the process whereby neurons are generated from neural stem cells. Even though most neuronal populations are generated during embryonic development, there is evidence in mammals that adult neurogenesis continues throughout life. This occurs in the subgranular zone (SGZ) of the dentate gyrus (DG) and the subventricular zone (SVZ) of the lateral ventricle, niches where radial glia-like stem cell pools continue to proliferate and generate progeny that differentiate into neurons [1, 2]. The neuroblasts generated in the rodent SVZ migrate through the rostral migratory stream (RMS) to the olfactory bulbs (OB). Neurogenesis can be influenced by many growth factors, signalling molecules and transcription factors regulating the stem cell niche [3–5]. The rate of neurogenesis can also be influenced by environmental factors including stress, physical excercise or environmental enrichment [6–8]. Notably, exogenous intervention with chemicals or drugs such as selective serotonin reuptake inhibitors (SSRIs) can also be used to target neurogenesis [9].

The SGZ is of particular importance in humans, as it is implicated not only in memory, learning and affective behaviours, but is also affected in diseases like Alzheimer’s disease (AD), epilepsy and schizophrenia [2]. While it is well-accepted that adult neurogenesis occurs in almost all mammals, the extent to which it occurs in adult humans has long been debated [10, 1, 11]. Some of the first evidence supporting adult neurogenesis in humans was found in patients who had been given bromodeoxyuridine (BrdU) to monitor their cancers [10]. These patients had BrdU+ cells in the SVZ and displayed BrdU+NeuN+ double-positive cells in the DG even past the age of 70, thus suggesting an active hippocampal neurogenic niche. Other reports followed, including carbon dating individual neurons in different areas of the adult brain which revealed that SVZ derived neurogenesis produces caudate nucleus neurons and that the SGZ daily produces many hundreds of newborn neurons throughout life [10, 12–15, 11]. In a series of carefully planned studies, which unlike many other human postmortem studies used rigorous and reproducible histology, neurogenesis was detected in healthy people into the 9^th^ decade of life but was decreased in AD and other neurological disorders [11, 16, 17]. This was supported by a recent single cell RNA sequencing (scRNAseq) study of the human brain that found imature DG granule cells throughout life as well as cells with stem cell like transcriptomes that were shown to be proliferative and neurogenic in vitro and in vivo [18].

Over forty-six million people worldwide are estimated to live with dementia [19]. AD is the most common form of dementia and accounts for 60-80% of all cases. It is a progressive and fatal neurodegenerative disease associated with age and includes symptoms such as memory loss, personality changes, loss of language and motor skills, and depression [20]. AD is characterised by the presence of amyloid-β (Aβ) and tau hyperphosphorylation in the form of senile plaques and neurofibrillary tangles (NFT), respectively [21]. The current therapeutic strategies, mainly based on acetylcholinesterase inhibition, are not disease modifying and only improve symptoms during a limited period of time [22]. Recently, the first disease modifying AD drug, aducanumab, has been approved by the U.S. Food and Drug administration (FDA). However, historically disease modifying therapies, notably monoclonal antibody treatments (targeting amyloid build-up) have failed to meet clinical endpoints. It is noteworthy that alterations in adult neurogenesis have been implicated in AD pathogenesis, not only because the DG is one of the first areas to be affected, but also because some of the key molecules underlying AD have important roles in the stem cell niche regulation [23]. Furthermore, some of the putative physiological functions of neurogenesis, such as pattern separation and spatial memory, and its link to depression when downregulated are also associated with AD [24, 25].

A more direct link between human AD and neurogenesis has been investigated by looking at newborn protein markers such as doublecortin (Dcx), microtubule associated protein 2 (Map2) or polysialylated nerve cell adhesion molecule (PSA-NCAM) by immunohistochemistry in post-mortem samples of AD patients compared to age-matched individuals. The same study that recently confirmed human hippocampal neurogenesis in neurologically healthy individuals also showed it significantly decreases in Alzheimer’s disease [11]. However, there is evidence that different phases of the disease and/or disease severity can differentially regulate adult neurogenesis. Many studies show decreases in adult neurogenesis in AD [26, 27, 11], whereas others claim no change [28] or increased numbers of new-born or immature neurons in the DG of AD patients [29]. Some of the discrepancy may be explained by post-mortem interval (PMI) differences, especially considering that Dcx is very susceptible to PMI and overfixation [28, 16], but also by the stage of disease progression. These studies collectively provide not only more evidence for human hippocampal neurogenesis but also suggest that declining neurogenesis in AD could be targeted to ameliorate the disease. What is clear is that increasing neurogenesis in both animals and humans using pharmacological intervention (i.e., SSRIs) or exercise can have beneficial effects, and given the potential positive ramifications of targeting stem cells, there has been more focus on stimulation of endogenous stem cell populations to galvanize improvements in patients. This could have large ramifications for future treatments in conditions where there are limited or none therapies available.

We reasoned that combining the well-established field of drug discovery with the rather newer concept of stem cell therapy is a tractable goal. We decided to use a phenotypic screen as many new drugs have been found with this approach [30]. Others have used a similar approach but pooled molecules and screened for in vivo increases in hippocampal neurogenesis [31]. In order to identify novel chemical matter modulating the complex biology behind the process of neurogenesis, we ran in vitro phenotypic screening using murine primary neural stem cells (NSC). Here we report that one of the hits, OXS-N1, has pro-neurogenic effects in both the hypocampus and the SVZ/OB of wild-type (WT) mice.

Moreover, behaviroral tests showed that treatment with OXS-N1 can lead to improvements in spatial memory in WT mice and an AD mouse model, supporting the idea that enhancing neurogenesis could improve behavioural symptoms in AD.

## Methods

### ADME and pharmacokinetic studies

All absorption, distribution, metabolism and excretion (ADME) studies were outsourced to Cyprotex, UK. This included performing plasma protein binding, permeability, multidrug resistance protein 1 (P-GP) efflux- Madin-Darby canine kidney (MDCK) studies, hepatocyte stability studies. Pharmacokinetic (Pk) studies which were performed by Cyprotex used male CD1 mice which were administered compound in 90% phosphate buffered saline (PBS), 10% dimethyl sulfoxide DMSO via intraperitoneal (IP) or gavage with n=3 for each time point.

Animals were then killed at 8 time points over 24 hrs and plasma and brain levels of compound measured using liquid chromatography–mass spectrometry (LC-MS). For excretion studies animals administered compound had their urine and faeces tested over a 24hr period by LCMS.

### Primary neural stem cell neurosphere culture

Primary neurospheres were obtained by separately culturing SVZ and SGZ cells isolated from postnatal day (P)3-6 C57BL/6 mice using 2-3 animals per culture. Animals were administered an overdose of pentobarbital followed by cervical dislocation and the brains removed and placed in a petri dish containing cold Hanks balanced salt solution (HBSS) (GiBCO 14025092). In order to access the stem cell niches, the brains were sliced in 500 μm coronal sections using a McIlwain tissue chopper. Sections containing the SVZ and SGZ in the DG were microdissected separately in HBSS using a dissection microscope (Leica) and stored in 15 mL tubes on ice. In order to avoid inclusion of SVZ cells in the SGZ, we specifically only included the DG and not the wall of the lateral ventricles. After collecting the tissue from all brains, 1 mL of 1 x accutase solution (A6964, Sigma-Aldrich) was added to the pooled tissue in each tube, gently mixed and incubated for 15 min at 37°C. Mechanical dissociation via trituration with a 200 µL Pipetteman was carried out after the incubation to obtain a single cell suspension, followed by addition of 12mL of NB-A+ (Neurobasal-A media 1X GiBCO-10888022 / 1X B27 GiBCO-17504 / 1X Glutamax GiBCO 35050061 / 100U/ml Penicillin-Streptomycin GiBCO 15140122) supplemented with human epidermal growth factor (hEGF) (20 ng/ml, Sigma E9644) and human fibroblast growth factor 2 (hFGF2) (20 ng/mL, R&D systems 233-FB-025) freshly added into the media. The single cell suspension was then plated in a 6 well plate (Corning 351146) and incubated at 37°C with 5% CO2. The day after, the media was changed, and the cells were left in the incubator, with a media exchange performed the following day. On day 6, the neurospheres were passaged using accutase, re-suspended in NB-A+ supplemented with hEGF & hFGF2 and plated in Ultra-Low Attachment cell culture T75 flasks (Corning CLS3814) at 5 x 10^4^ cells/mL. The following passages were carried out every 5 days and cells were always used for experiments as tertiary neurospheres, i.e., after the second passage.

### Neurosphere Differentiation Assay

For compound screening, 96 or 384 well plates were coated for adherence with 15 µg/mL poly-ornithine (Sigma P3655-50MG) in PBS (ThermoFisher 10010015) overnight at 4°C, then washed once with PBS and incubated with 10 µg/mL of laminin (Sigma L2020) in PBS for at least 1 hr at 37°C. Cells were dissociated with accutase, re-suspended in NB-A+ without growth factors and seeded as monolayers in 96 well plates (25,000 cells/well) or in 384 well plates (7,500 cells/well). The plates were incubated for 1 hr for the cells to settle down before adding the test compounds diluted in NB-A+. The compounds used in all the experiments were diluted in 100% DMSO to generate a 10 mM stock, and the final concentration of DMSO administered to the cells was 0.1%. After compound addition, the cells were left in the incubator to differentiate for 6 days without any media change, washed once with PBS and then fixed in 4% paraformaldehyde (PFA, Santa Cruz Biotechnology, sc- 281692), with rocking for 15 minutes at room temperature (RT). Subsequently, the cells were immunostained and analysed.

### Immunocytochemistry

After fixation, cells were washed once with PBS for 10 min followed by a blocking-permeabilization step for 1 hr at RT in blocking buffer (PBS/ 5% donkey serum Bio-Rad C06SB / 0.1% Triton Sigma X100). Primary antibodies diluted in blocking buffer were then added to the cells (Table 1) and incubated at 4°C overnight on a rocking shaker. The following day, the cells were washed once in PBS for 10 min and incubated with the appropriate Alexa Fluor secondary antibodies (1:500, Invitrogen) for 1 hr at RT. The cells were then washed in PBS for 10 min, incubated with DAPI (1 μg/mL, Sigma D9542) for 5 min at RT to label cell nuclei and washed again for 10 min. Plates were imaged on a Perkin Elmer Operetta or EVOS FL auto, acquiring a minimum of 4 images per well, and analysed using ImageJ and Cell Profiler excluding dead cells. Ommision of primary antibodies was used as a negative control and resulted in neglibeable background stain. Cells were counted by an observer blinded to drug or vehicle exposure, and randomly generated identification codes were only revealed post-quantification.

**Table 1.** Primary antibodies used *in vivo*.

| Antigen | Dilution | Host | Manufacturer | Product Cat. No. |
| --- | --- | --- | --- | --- |
| BrdU | 1:400 | Sheep | Abcam | Ab1893 |
| BrdU | 1:400 | Rat | Novusbio | NB500-169 |
| Dcx | 1:100 | Goat | Santa Cruz | Sc-8066 |
| GFAP | 1:400 | Rat | Life Technologies | 13-0300 |
| NeuN |  |  |  |  |
| pHH3 | 1:400 | Rabbit | Abcam | Ab47297 |
| Ki67 | 1:400 | Rat | Abcam | Ab15580 |

### Animals

P3-6 C57BL/6 mice (males and females) obtained from the University of Oxford Biomedical Science service were used for all *in vitro* cultures. Female C57BL/6J mice obtained from Charles River Laboratories were used at 3 months of age for wild type (WT) *in vivo* studies. For pharmacokinetic (PK) studies female CD1 animals at 3 months of age were used.

B6.Cg-*Zbtb20^Tg(PDGFB-APPSwInd)20Lms^*/2Mmjax (J20) transgenic mice were obtained from JAX (Jackson Laboratories, ME) and male heterozygous animals were bred with female WT of the same background strain. Animals were genotyped at P21 by Transnetyx, using ear clips. For compound administration studies in J20s both male and females between the ages of 6.5-11.5 months old were used. Animals were maintained in individually ventilated cages on a 12-h light/dark cycle with food and water provided *ad libitum*. Procedures were performed with University of Oxford Research Ethics Committee approval in accordance with the Animals (Scientific Procedures) Act of 1986 (UK). All efforts were made to minimize animal suffering and distress.

### OXS-N1 administration

WT or J20 animals were treated with 25 mg/kg of OXS-N1 (5% DMSO, PBS) or vehicle (5% DMSO, PBS) 3 times a day via IP injections or gavage (PO) administration for 7 or 21 days. Animals were administered 0.8 mg/mL BrdU in the drinking water, supplemented with 1% sucrose for either the duration of dosing (all WT studies) or the final 7 days during 21-day dosing (J20s). Animals which received 7 days of OXS-N1 had a 2-week washout period and animals treated for 21 days had a 3-week washout period.

### Behavioural testing

#### Spatial Novelty Preference in the Y Maze

A Y maze made of Perspex with 3 arms of 35cm in length, 8cm wide and 10cm high with an angle of 120° between each arm, with extra-maze visual cues places around the maze, was used. All apparati were cleaned with 70% ethanol between animals and lighting was set at 30 ± 5 lux. Dirty bedding from animals of the same sex (but not included in the study) was mixed in with two thirds clean bedding and lined the floor of the maze. Animals were allowed to acclimatise to the room for 1 hr before the test. Initially, the animal was allowed to explore two arms of the maze (the start and familiar arms) with the third alternating arm blocked off. The animal was allowed to explore for 5 minutes. Animals were then removed from the maze for 2 minutes, the bedding mixed to avoid olfactory cues and the third, novel arm exposed. The animals were placed back in the maze and the time spent in the novel arm was recorded using AnyMaze software. Memory performance was expressed as the time spent in the novel arm as a percentage of the time spent in the novel arm and the familiar arm combined (i.e. chance performance is 50%).

#### Locomotion

Locomotor activity (LMA) boxes were used which comprised a transparent Plexiglas box (48 × 27 × 21 cm, *L* × *W* × *H*, Photo Beam Activity Hardware and Software, Open Field San Diego Instruments), filled with clean sawdust and a plexiglas lid, with perforations. Animals were allowed to explore freely for 1 hr. LMA was recorded based on the number of horizontal beam breaks made by the animals. All apparati were cleaned with 70% ethanol between animals and lighting was set at 30 ± 5 lux.

### Immunohistochemistry

Animals were overdosed with euthatal and transcardially perfused with cold PBS and 4% PFA. Brains were removed and post fixed in 4% PFA overnight and then cryoprotected in 30% sucrose in 0.1M phosphate buffer for 3 days. Brains were frozen in dry ice then sectioned into 30 μm coronal sections using a freezing microtome (Leica).

Immunohistochemistry was then performed as previously described [32]. Primary antibodies used are shown in Table 1 and Alexa Fluor secondary antibodies were used (1:500, Invitrogen). Antigen retrieval for BrdU staining was performed with 1M HCl at 37°C for 1 hr. Images were taken on a Zeiss LSM 710 laser scanning confocal microscope with 1µM Z- stacks and tile scans as appropriate, or on an automated EVOS FL Auto2 (Invitrogen) cell and slide imaging system and quantified using ImageJ. For each immunohistochemistry we omitted the primary antibody as a negative control to reveal any background staining and only counted cells that were clearly more fluorescent than the background.

### Transcriptomics

We carried out bulk RNAseq on SVZ and SGZ neurosphere cells exposed for 3 days to OXS- N1. Altogether there are 16 samples, 8 from each region (SVZ, DG), 4 for each state (prolif, differentiation) and 2 for each condition (treated, control). Differential gene expression analysis was caried out with DESeq2 package. Gene Ontology enrichment was done using the clusterProfiler package. All transcriptomics analyses were performed in R.

In other experiments we carried out bulk transcriptomics post-OXS-N1 on human induced pluriptential stem cell (hIPSC) derived neurons. hiPSC cortical neuronal maturation was accelerated by overexpression of NGN2 as in [33] and OXS-N1 was administered for 2 days. Cases 010 (female, 17.8 yrs) and 014 (male, 14.9 years) each had two lines tested.

### Statistics

Data were expressed as mean ± standard error of mean (SEM). All *in vitro* experiments were performed using at least 3 technical and 3 biological replicates. All *in vivo* data had 8 animals per group for behaviour and subsequent immunohistochemical analysis, unless otherwise stated. Data analysis was performed using GraphPad Prism 7.05. All data were analysed initially using a Brown-Forsythe’s test to test for equality of variances, and Shapiro-Wilk test for normality. If data were normally distributed either an unpaired Student’s t-test (for 2 groups, equal variance unless otherwise stated) or one way ANOVA (2+ groups) followed by post-hoc Tukey’s t-test was performed. If data were not normally distributed then a Mann-Whitney test (for 2 groups) or a Kruskal*–*Wallis test (2+ groups) were used, followed by Mann-Whitney test post-hoc. Values given are (mean ± s.e.m.). *p<0.05, **p<0.01, ***p<0.001.

## Results

### An *in silico* and phenotypic screen using DG and SVZ derived NSCs identified small organic molecules that increased neurogenesis

1,080 compounds were screened from a Biofocus library based on *in silico* pharmacophore modeling (Figure 1A, Figure S1, and Supplementary Methods). This was achieved using a virtual screening cascade (Figure 1A) to select compounds similar to known neurogenic compounds. This included similarity screens using Knime. Figures 1A and S1, represent the screening cascade adopted with input from compounds identified through the in-silico modelling. We sought to avoid over-activating and thus depleting stem cells and therefore, compounds were screened during the differentiation phase of the neurosphere assay at one concentration of 10 µM in SVZ derived NSCs.

**Figure 1:**
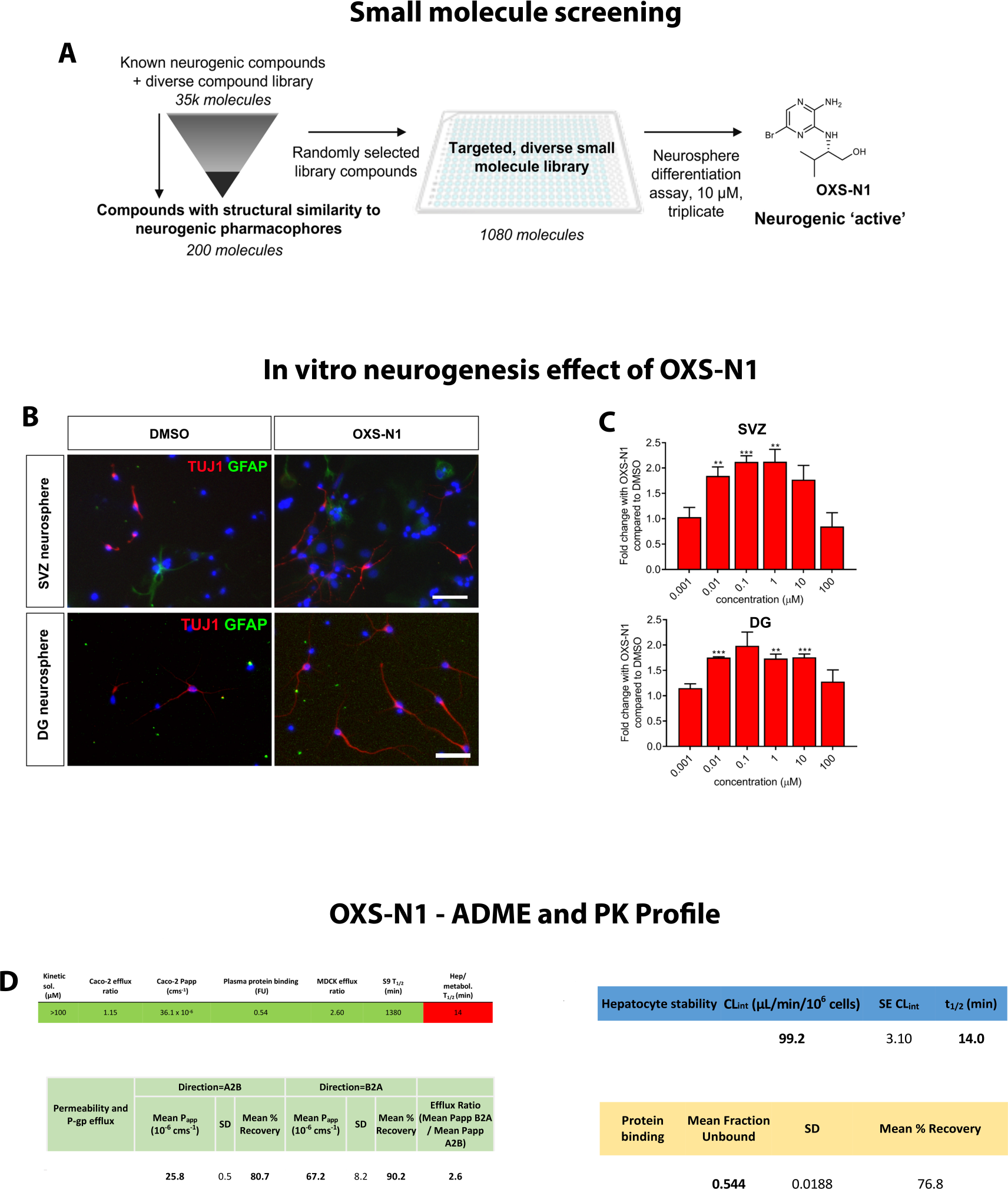
In vitro drug discovery pipeline. (A) Schematic of *in vitro* screening strategy optimised to identify compounds capable of enhancing neurogenesis in a physiological primary cell culture model of adult neurogenesis (neurosphere assay). (B) Images of cells from dentate gyrus (DG) or subventricular zone (SVZ) neurospheres differentiated for 6 days in vitro (DIV) in the presence of OXS-N1 or vehicle (DMSO). Cultures were immunofluorescently labelled for TUJ1 (neurons, red) and GFAP (astrocytes, green) and DAPI (cell nuclei, blue). Scale bar = 50μm. (C) Cultures were assessed for the percentage of neurons present. Concentration response curve for DG and SVZ derived cells, quantified as fold-change in percentage of neurons compared to DMSO (mean ± s.e.m.). Statistically significant increases compared to DMSO are indicated (* p<0.05 ** p<0.01 ***p<0.001). (D) absorption, distribution, metabolism, excretion rates for OXS-N1.

Immunocytochemistry on both SVZ and DG neurospheres was then performed using neuronal and glial markers (Figure 1B) and the percentage of Tuj1+ young neurons was calculated. Compounds that resulted in significantly greater neurogenesis than DMSO controls were then further screened in SVZ and DG neurospheres, with concentration response assays of resynthesised chemical matter (Figure S1). Concurrently, evaluation of absorption, distribution, metabolism and excretion (ADME) was performed on compounds which significantly increased the neuronal population *in vitro* (Figure S1). Efficacious compounds were then administered IP or PO into male mice and brains collected at 8 time points over 24 hrs to measure brain and plasma concentrations. Compounds which demonstrated reasonable systemic exposure in both blood and the brain were then be tested in WT mice and those that showed positive effects on neurogenesis were used in an AD mouse model.

From the 1,080 compounds that were screened in the phenotypic assays 30 significantly increased neurogenesis in SVZ and or on SGZ neurospheres or in both (2.8% hit rate), and these were stratified into 5 chemical series (Figure 1A). Of these compounds, 5 were found to produce a significant increase in neurons compared to controls *in vitro* and to also have desirable ADME properties. PK analysis revealed that only 2 compounds were likely capable of penetrating the blood brain barrier (BBB) after systemic exposure. One of these compounds, designated OXS-N1, significantly increased the proportion of neurons in both DG and SVZ derived cultures after 6 days of treatment and was comparable in efficacy with a positive control retinoic acid (RA) treatment (Figure 1B,C). ADME analysis revealed that OXS-N1 was actively effluxed in a *MDCK* efflux assay and had an efflux ratio of 2.6 (Figure 1D), indicative that it may cross the blood brain barrier. It also showed a clearance level of 99.2 minutes in a liver microsomal hepatocyte stability assay (Figure 1F), suggesting that it would be cleared rapidly upon systemic administration.

### Intraperitoneal administration of OXS-N1 increases SGZ neurogenesis in WT mice

To investigate whether OXS-N1, a compound capable of increasing neuronal populations *in vitro*, was able to recapitulate this effect *in vivo,* we assessed its effect in WT mice. Initially a PK study was performed, whereby animals were intraperitoneally injected with 5 mg/kg of OXS-N1 and killed at intervals to assess plasma and brain exposure. OXS-N1 was detected at up to 1 hr in the plasma and brain of injected mice (Figure 3A).

Although this demonstrated reasonable exposure, we increased the concentration of future dosing to 25 mg/kg to ensure that we potential reached therapeutic levels for a prolonged time. The administration paradigm (Figure 2B) consisted of one week of compound administration to target the neurogenic niches followed by a two-week washout period to enable functional integration of any newborn neurons and for examination of behavioural effects on hippocampus-dependent tests. Three-month old, female, C57BL/6J mice were used (n=8/group) and animals were dosed IP three-times daily (TID) for 7 days with either OXS-N1 or vehicle, and BrdU given concurrently via the drinking water (Figure 2B).

**Figure 2:**
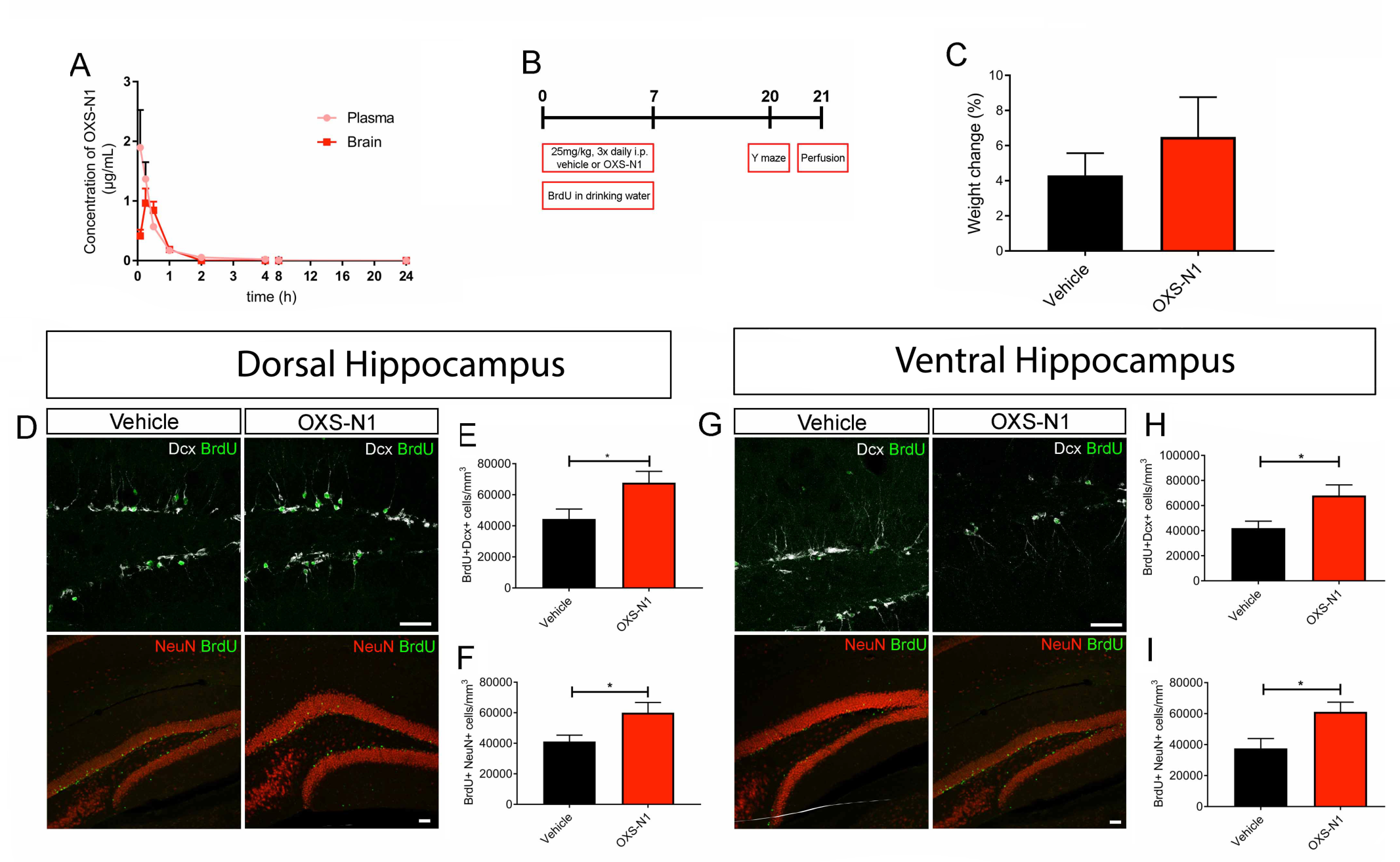
One-week IP administration of OXS-N1 increases adult neurogenesis in wild- type mice. (A) Concentration of OXS-N1 in plasma and brain at various time points after a single IP dose of 5 mg/kg. (B) Experimental timeline for study design for one-week, three- times daily (TID) IP administration of OXS-N1 to wild-type female mice (12 weeks of age at the start of the study). (C) Change in body weight of animals between first day and last day of dosing. (D) Maximum intensity projection images of confocal z-stacks taken in the dorsal dentate gyrus of brains immunofluorescently labelled for BrdU and Dcx or BrdU and NeuN. (E) Quantification of the density of BrdU+Dcx+ double-labelled new-born neurons in the dorsal dentate gyrus. (F) Quantification of the density of BrdU+NeuN+ double-labelled mature neurons in the dorsal dentate gyrus. (G) Maximum intensity projection images of confocal z-stacks taken in the ventral dentate gyrus of brains labelled for either BrdU and Dcx or BrdU and NeuN. (H) Quantification of the density of BrdU+Dcx+ double-labelled new-born neurons in the posterior dentate gyrus. (I) Quantification of the density of BrdU+NeuN+ double-labelled mature neurons in the ventral dentate gyrus. Dcx: doublecortin, NeuN: Neuronal nuclei, BrdU: Bromodeoxyuridine. Scale bars = 50 μm.

Extensive monitoring was performed to ensure that there was no obvious toxicity, as this compound had never been administered *in vivo* to our knowledge. Mice were monitored daily for potential adverse effects following a checklist of readouts including, but not limited to, discharge, tremors, limb dragging, gait, hypermetria and proprioception. Neither these, nor any other adverse effects were observed during the study. Animal weights were monitored and although the OXS-N1-treated mice gained somewhat more weight than the controls, the difference was not statistically significant (t-test p= 0.41; Figure 2C).

We first examined the dorsal DG which is involved with spatial memory (Figure 2D-F) [34]. To explore the effect of OXS-N1 on neurogenesis *in vivo*, we used immunohistochemistry to detect BrdU+Dcx+ double-labelled cells in the DG (Figure 2E). Doublecortin (Dcx) is a microtubule-associated protein necessary for SVZ neuroblast migration [35] and expressed for approximately two weeks after SVZ and SGZ neuroblasts are born [36, 37]. NeuN begins to be expressed by SVZ and SGZ neuroblasts approximately two weeks after they are born [38]. There were significantly more BrdU+Dcx+ cells in the dorsal DG of animals treated with OXS-N1 compared to vehicle (Figure 2E) (44,460 cells/mm^3^ ± 6,387 (vehicle) vs. 67,784 cells/mm^3^ ± 7,322 (OXS-N1); t-test p=0.0308. To assess whether the increase in newborn neurons after OXS-N1 administration is reflected in cells past the initial stages of maturation, we examined the mature neuronal population double-labelled for BrdU+NeuN+ in the DG (Figure 2D). We also found a significant increase in the number of BrdU+NeuN+ double-labelled cells in the granule layer of the dorsal DG of OXS-N1 treated animals (Figure 2F) (39,876 cells/mm^3^ ± 3,490 (vehicle) vs. 60,040 cells/mm^3^ ± 6,739 (OXS-N1); Welch’s t-test p=0.0231)

We next examined the ventral DG (Figure 2G-I) which has functions in anxiety [39–41, 34]. In the ventral DG there were also more BrdU+Dcx+ double positive cells: 42,024cells/mm^3^±5,592 (vehicle) vs. 68,019cells/mm^3^±8,495 (OXS-N1); t-test p=0.0229) (Figure 2H). Furthermore, the ventral DG contained more BrdU+NeuN+ cells in the drug administration group: 37,277cells/mm^3^±6,145 (vehicle) vs. 58,845cells/mm^3^±5,913 (OXS- N1); Welch’s t-test p=0.0241) (Figure 2I). This suggests survival of the additional newborn neurons in the DG niche, or increased differentiation into mature neurons. Notably, neurogenesis was significantly increased in OXS-N1 treated mice in both the dorsal and the ventral portions of the DG which are known to have different functions [42, 39, 43, 40, 41, 44–46].

Glial fibrillary acidic protein (GFAP) is an intermediate filament marker of the astrocytes and of the astrocyte-like stem cells of the DG. To further explore the action of OXS-N1 and determine if stem cells became activated by OXS-N1 administration, we immunostained for BrdU+GFAP+ cells. Significantly more BrdU+GFAP+ double-labelled cells were detected in the ventral DG of OXS-N1 treaed mice: (46,024 cells/mm^3^ ± 12,680, n=8) compared to control mice (19,299 cells/mm^3^ ± 4,622 p=0.0004, n=6) (data not shown). This effect was not detected in the dorsal DG (4,291 cells/mm^3^ ± 3,340 (vehicle) vs. 4,068 cells/mm^3^ ± 3,212 (OXS-N1, p=0.89, n=7 brains for vehicle and n=8 brains for OXS-N1). Most proliferation in the SGZ is not of the GFAP+ NSC but of cells further advanced in the lineage; intermediate progenitors and neuroblasts. To determine overall rates of proliferation at the end of the two-week washout, we quantified the number of pHH3+Ki67+ cells (actively proliferating cells at the time of fixation two weeks after the end of compound administration). Neither single nor double-labelled proliferative cells were significantly different in number between OXS-N1 and control mice in the dorsal and ventral DG (Figure S2A). We next determined if OXS-N1 affects proliferation in the SVZ, by quantifying the number of pHH3+Ki67+ cells (Figure S2B). There were also no statistical differences in the number of BrdU+NeuN+ mature neurons in the olfactory bulb (n=8 brains for vehicle and n=6 brains of OXS-N1) (data not shown).

Taken together these data shows an increase in neurogenesis in the dentate gyrus upon IP injections of OXS-N1 for one week as evidenced by an increase in the neuroblast and mature neuronal populations.

### Oral administration of OXS-N1 increases SGZ and OB neurogenesis in WT mice

To establish a more refined mode of delivery, we performed a pharmacokinetic (PK) study using oral administration via gavage. Animals were dosed with 25 mg/kg OXS-N1 and killed at different time points over 24 hrs, after which brain and plasma compound concentration was measured (Figure 3A). The brain concentration after 30 min was roughly comparable to that after IP dosing, when considering that the PK curve for IP was generated at 5mg/kg and that for p.o. dosing at 25mg/kg. OXS-N1 reached µM blood and brain levels upon PO administration using oral gavage. Thus, oral and IP routes of administration should be both valid and comparable. 3-month-old female C57BL/6J mice (n=8/group) were administered with either OXS-N1 or vehicle PO TID for either 7 or 21 days (Figure 3B), with BrdU administered concurrently by dosing via the drinking water. Animals were closely monitored in the 7- and 21-day study, and no adverse effects were observed. Nor were there significant weight differences between vehicle and OXS-N1 dosed mice for either length of administration (Figure 3C).

**Figure 3:**
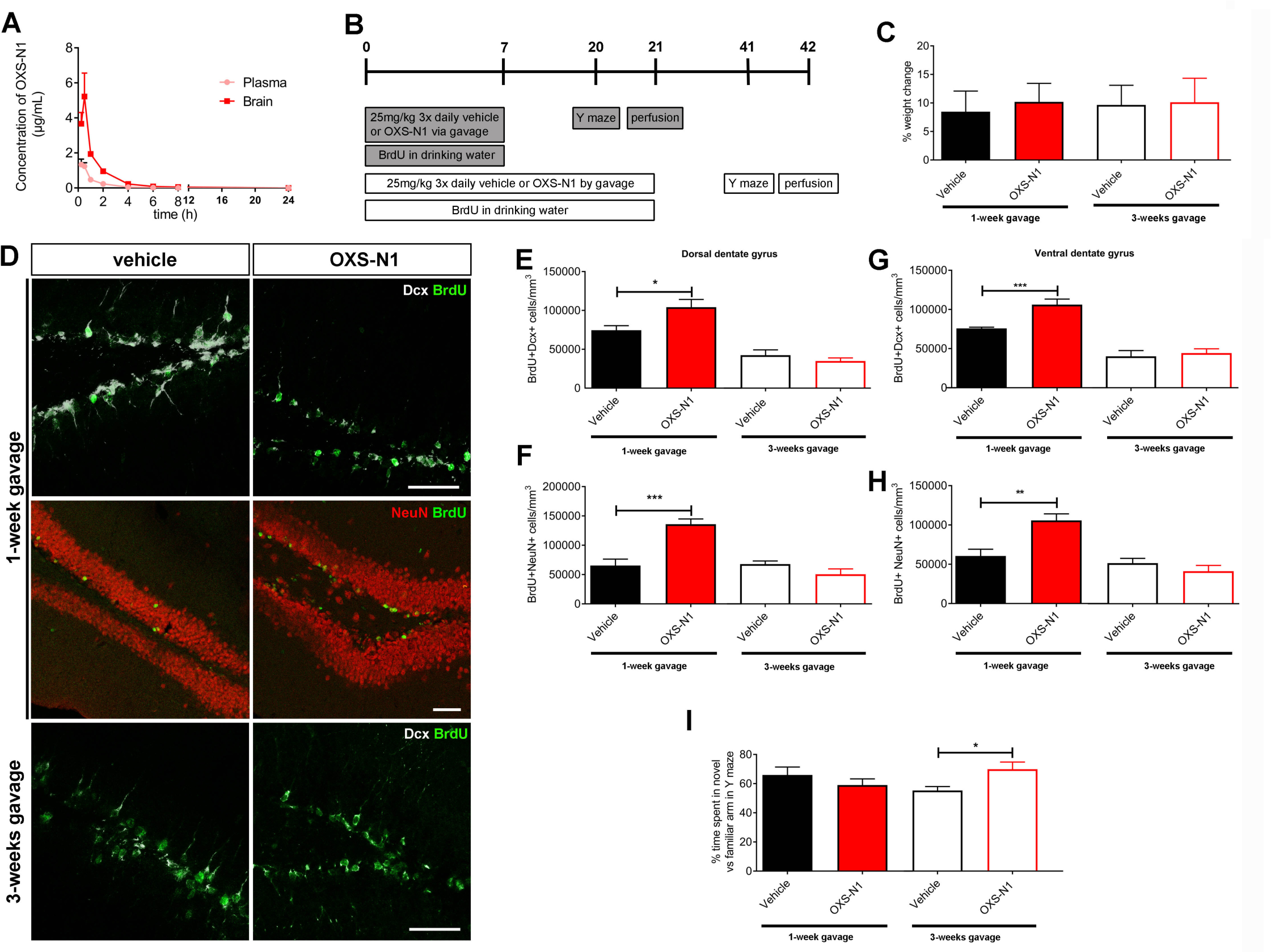
One-week oral administration of OXS-N1 increases adult neurogenesis in wild-type mice, and three-weeks administration improves memory. (A) Concentration of OXS-N1 in plasma and brain at various time points over 24 hrs after a single p.o. (gavage) dose of 25mg/kg. (B) Experimental time-line of study, one- and three-weeks of TID p.o. gavage administration of OXS-N1 to wild-type female mice (12 weeks of age at the start of the study). (C) Change in body weight of animals between first day and last day of experiment. There were no significant differences between drug and vehicle dosed animals. (D) Tissue sections were immunofluorescently labelled with antibodies against Dcx and BrdU or NeuN and BrdU to determine whether one week and three weeks of OXS-N1 administration can influence adult neurogenesis in this niche. Maximum intensity projection images of confocal z-stacks. (E) Quantification of the density of BrdU+Dcx+ double-labelled new-born neurons in the dorsal dentate gyrus. There were significantly more BrdU+Dcx+ cells after 1-week of p.o. dosing with OXS-N1 compared to vehicle controls (p=0.026, unpaired t-test), but not after three weeks of dosing. (F) Quantification of the density of BrdU+NeuN+ double-labelled mature neurons in the dorsal dentate gyrus. The density of BrdU+NeuN+ double-labelled mature neurons was significantly higher in the group treated with OXS-N1 for one week, compared to the vehicle group (p=0.0002, unpaired t-test), but not significantly different after three weeks of dosing. (G) quantification of the density of BrdU+Dcx+ double-labelled new-born neurons in the ventral dentate gyrus. There were significantly more BrdU+Dcx+ cells after 1-week of p.o. dosing with OXS-N1 compared to vehicle controls (p=0.0002, Mann-Whitney test). (H) quantification of the density of BrdU+NeuN+ double-labelled mature neurons in the ventral dentate gyrus. The density of BrdU+NeuN+ double-labelled mature neurons was significantly higher in the group treated with OXS-N1 for one week, compared to the vehicle group (p=0.0047, Mann-Whitney test), with no obvious difference following three weeks of treatment. (I) Animals were tested in a Y-maze 13 days after the end of drug dosing to determine whether OXS-N1 has a cognitive effect after one or three weeks of oral administration. Memory performance was expressed as the time spent in the novel arm as a percentage of the time spent in the novel arm and the familiar arm combined (i.e. chance performance is 50%)., Memory performance was significantly higher in the group treated with OXS-N1 for three weeks (p=0.0217, unpaired t- test). Dcx: doublecortin, NeuN: Neuronal nuclei, BrdU: bromodeoxyuridine. Scale bars = 50 μm.

As before, we examined the BrdU+Dcx+ double-labelled population of cells in the DG, to determine the effect of orally administered OXS-N1 on neurogenesis *in vivo* (Figure 3D). Immunofluorescence analysis of the hippocampal stem cell niche of brains from animals receiving one week but not three weeks of oral OXS-N1 revealed a significant increase in the BrdU+Dcx+ neuroblast population of newly generated neurons (Figure 3E,G). The differences between the vehicle and the OXS-N1 group treated for one week were statistically significant in the dorsal dentate gyrus (74,549 cells/mm^3^ ± 5,925 (vehicle) vs. 104, 241cells/mm^3^ ± 9,960 (OXS-N1) (n=8/group); p=0.0226 t-test). This was also the case in the ventral dentate gyrus (75,798 cells/mm^3^ ± 1,611 (vehicle, n=8) vs. 106,126 cells/mm^3^ ± 7,155 (OXS-N1, n=7); p=0.0002 Mann-Whitney test).

Overall, the density of BrdU+Dcx+ double-labelled cells appeared to be higher in the one-week group compared to the three-week group, in both untreated and treated cohorts. For example, in control brains there were 74,549 cells/mm^3^ ± 5,925, (vehicle 1-week, dorsal dentate gyrus) vs. 42,448 cells/mm^3^ ± 6,846 (vehicle 3-week, dorsal dentate gyrus). BrdU was administered throughout the period of drug dosing, but the wash-out period was one week longer after 3-weeks of dosing, which may suggest that more cells that picked up the BrdU label had progressed through differentiation, therefore downregulating expression of Dcx.

We therefore assessed the number of neurons born during the experiment that matured as indicated by BrdU+NeuN+ double labelling. The density of BrdU+NeuN+ mature neurons was much higher after one week of orally administered OXS-N1, compared to controls in both the dorsal (65,367 cells/mm^3^ ± 10,974 (vehicle, n=8) vs. 135,936 cells/mm^3^ ± 9,034 (OXS-N1, n=8); p=0.0002 t-test) and ventral dentate gyrus (60,502 cells/mm^3^ ± 8,720 (vehicle, n=8) vs. 105,765 cells/mm^3^ ± 8,453 (OXS-N1, n=8); p=0.0047 Mann-Whitney test) (Figure 3F,H). In contrast, after 3-weeks of treatment there was no significant difference in the density of BrdU+NeuN+ double-labelled cells between vehicle and orally administered OXS-N1 animals (Figure 3F,H). The overall density of these cells was comparable between animals treated with vehicle for 1-week and those treated with either vehicle or drug for 3 weeks. We next examined the activated stem cell population in the neurogenic compartment of the DG by counting the number of BrdU+GFAP+ label retaining cells in the SGZ. There were no significant differences in the density of BrdU+GFAP+ stem cells in the SGZ between vehicle and orally administered OXS-N1 groups, irrespective of length of administration (Figure S3).

Animals were tested in a novelty Y maze at the end of the respective washout periods for both dosing schedules (Figure 3B,I). Animals which were administered either vehicle or compound for 1 week did not show any differences in the proportion of time spent in the familiar and novel arm. However, 3 weeks of administration improved the short-term memory performance in OXS-N1 treated animals, which spent an average of 70% of their time in the novel arm suggesting enhanced hippocampal-dependent short-term memory (Figure 3I).

We next assessed SVZ neurogenesis in the one-week and three-week gavage groups with immunohistochemistry (Figure S4). There were no significant differences in the number of BrdU+Dcx+ cells in the SVZ of vehicle or OXS-N1 mice, after 1 week of gavage, nor after 3 weeks (Figure S4).

In contrast, in the OB core, the density of BrdU+Dcx+ neuroblasts was significantly greater after three weeks of oral OXS-N1 administration (Figure 4C): 23,399 cells/mm^3^ ± 2,804 (vehicle, n=7) vs. 38,655 cells/mm^3^ ± 4,656 (OXS-N1, n=8), t-test p=0.018). However, there was no significant difference at one week of oral gavage (38,056 cells/mm^3^ ± 3,905 (vehicle, n=7) vs. 38,415 cells/mm^3^ ± 4,834 (OXS-N1, n=7) (data not shown). Also, after three weeks of OXS-N1 oral administration, the density of mature neurons (BrdU+NeuN+) was significantly increased in the OB granular layer, (Figure 4E) (4,309 cells/mm^3^ ± 1,748 (vehicle, n=7) vs. 7,857 cells/mm^3^ ± 2,461 (OXS-N1, n=8), t-test p=0.0074). But this effect was not seen at one week of oral administration (data not shown). These 2 findings correlate since Dcx+ neuroblasts reaching the OB core from the RMS migrate and integrate into the granule cell layer where they become NeuN+ OB interneurons. Thus, a prolonged increase in the immature neurons could subsequently lead to increased integration of these more mature neurons at their destination. The data also suggest that three weeks of drug administration as well as a longer wash-out period combined to give enough time for neurons generated in the SVZ during the BrdU exposure to migrate into the OB.

**Figure 4:**
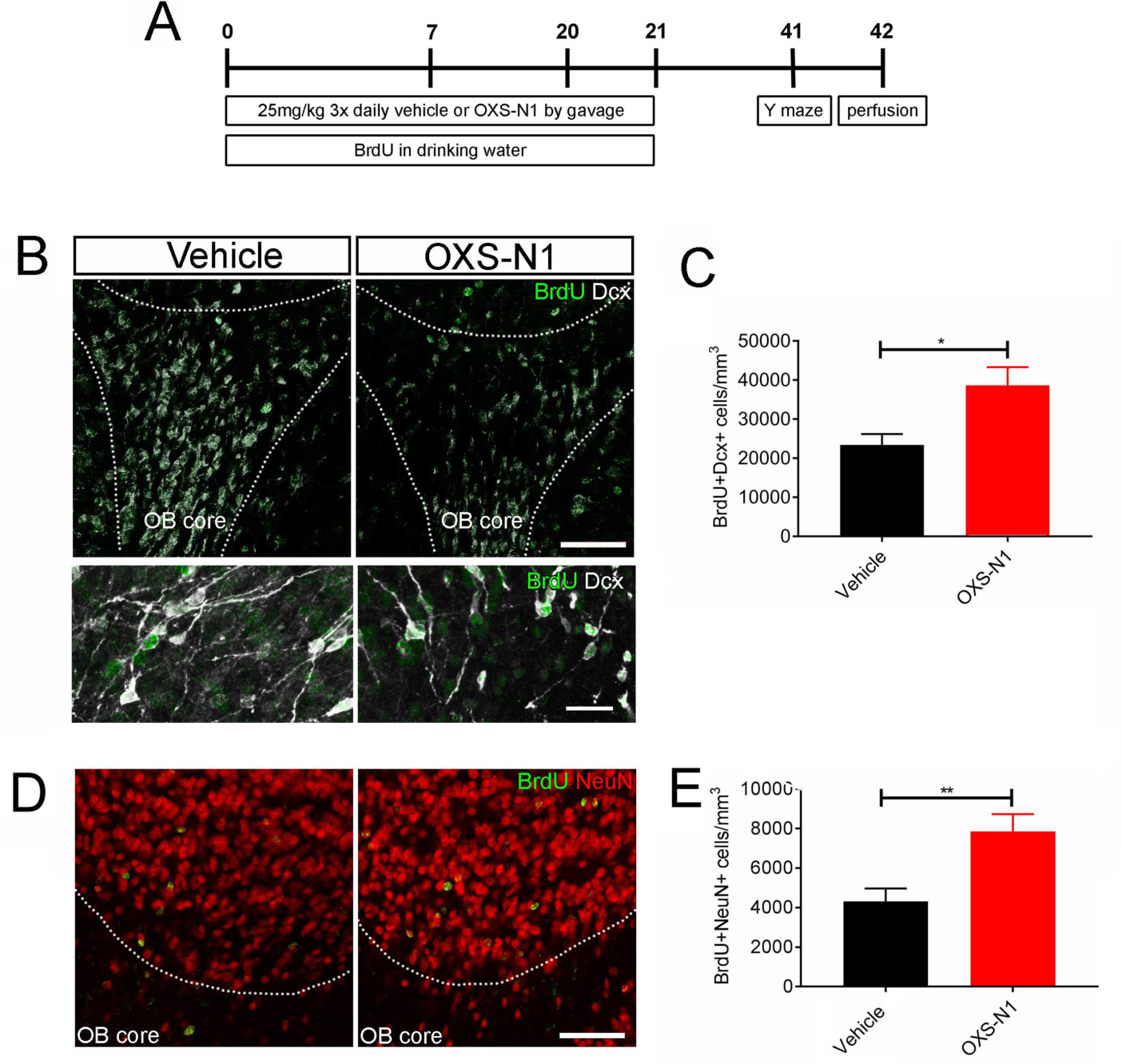
Three-week oral administration of OXS-N1 increases the density of new-born neurons in the olfactory bulb of wild-type mice. (A) Outline of experimental design. (B,D) Maximum intensity projection images of confocal z-stacks taken in the olfactory bulb of brains immunofluorescently labelled for either BrdU and Dcx or BrdU and NeuN. (C) Quantification of the density of BrdU+Dcx+ double-labelled new-born neurons in the core of the olfactory bulb. There was a significantly higher density of BrdU+Dcx+ double-labelled new-born neurons in the OXS-N1 treated animals compared to controls. (E) Quantification of the density of BrdU+NeuN+ double-labelled mature neurons in the granular layer of the olfactory bulb. The density of BrdU+NeuN+ double-labelled mature neurons was significantly higher in the OXS-N1 treated animals compared to the vehicle group (p=0.0188, unpaired t-test). Abbreviations: Dcx: doublecortin, NeuN: neuronal nuclei, BrdU: bromodeoxyuridine. Scale bars = 50μm.

### Oral administration of OXS-N1 increases neurogenesis and improves behavioural deficits in aged J20 mice

Heterozygous PDGF-APPSw,Ind (J20) mice, a genetic model of Alzheimer’s disease, reportedly have disrupted SGZ and SVZ neurogenesis from as early as 3 months and through 12 months of age [47]. OXS-N1 was administered to heterozygous J20 animals for three weeks via gavage (Figure 5A). BrdU was given via drinking water for the final week of drug administration, after which there was a three-week wash-out period. During this phase locomoter and novelty Y-maze behavioural tests were performed (Figure 5A). This length of washout was selected to allow newborn neurons to functionally integrate and contribute to behavioural results. The average age of the vehicle group (n=8) was 50.5±1.7 weeks, and for the compound group (n=8) was 50.7±2.7weeks. We used both sexes, as evenly distributed as possible (vehicle: 5 females, 3 males; OXS-N1: 6 females, 2 males).

**Figure 5:**
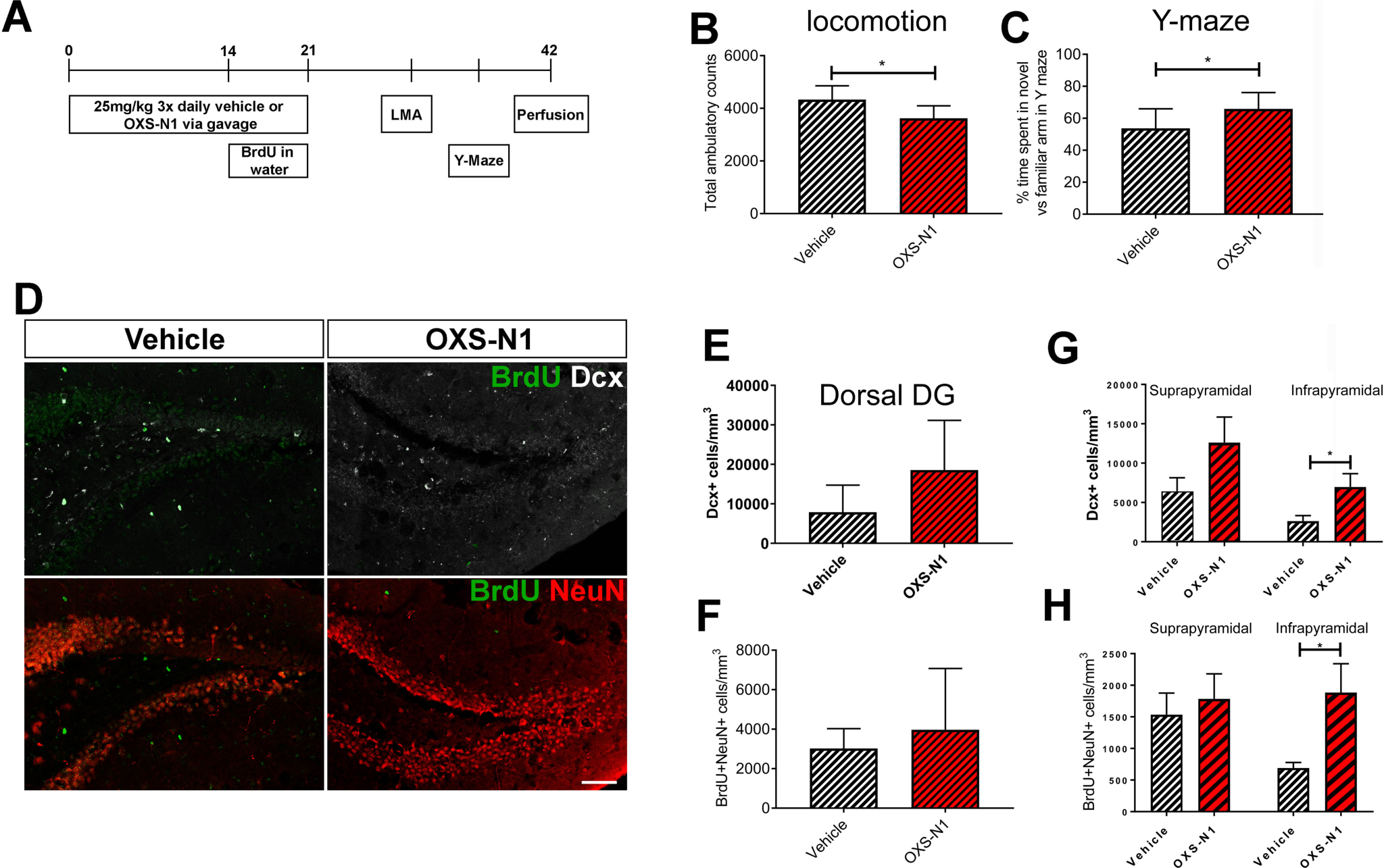
Three-week oral OXS-N1 improves behavioural deficits and increases neurogenesis in J20 mice. (A) Experimental timeline for study design three-week p.o. administration of OXS-N1 to one-year old J20 mice. Gavage was TID. (B) Total ambulatory movement counts were reduced in compound-treated J20s compared to vehicle controls (p=0.01, unpaired t-test). (C) Animals were tested in a Y-maze three weeks after the end of drug dosing. Memory performance was expressed as the time spent in the novel arm as a percentage of the time spent in the novel arm and the familiar arm combined (i.e. chance performance is 50%). Percentage of time spent in the novel arm was significantly higher in the group treated with OXS-N1 for three weeks (p=0.04, unpaired t-test). (D) Tissue sections from the dentate gyrus immunofluorescently labelled with antibodies against BrdU and Dcx or BrdU and NeuN. Maximum intensity projection images of confocal z-stacks taken in the dorsal dentate gyrus of brains. (E) Quantification of the density of Dcx+ new-born neurons in the dorsal dentate gyrus. (F) Quantification of the density of BrdU+NeuN+ double-labelled mature neurons in the dorsal dentate gyrus. (G) Quantification of Dcx+ cells in the suprapyramidal and infrapyramidal blades of the dorsal dentate gyrus showed that OXS-N1 significantly increased the number of Dcx+ cells in the infrapyramidal blade (p=0.01, unpaired t-test). (H) Quantification of BrdU+NeuN+ cells in the suprapyramidal and infrapyramidal blades of the dorsal dentate guyrus showed that OXS-N1 significantly increased the number of BrdU+NeuN+ cells in the infrapyramidal blade (p=0.01, unpaired t- test). Scale bar = 50 µm. LMA: locomotor activity, Dcx: doublecortin, NeuN: neuronal nuclei, BrdU: bromodeoxyuridine.

OXS-N1 treated J20s exhibited significantly reduced activity compared to vehicle treated J20 mice, as revealed by total ambulatory counts in a locomotor box for 1 hr (Figure 5B) (p=0.0137). This result indicates a potential correction of the well known J20 hyperactivity phenotype [48–50]. In the novelty Y maze test, compound-treated animals showed a significantly greater proporation of time in alternation to the novel arm (Figure 5C). Vehicle treated J20 animals (n=8) spent 54% ± 12 of time in the novel arm, whereas OXS-N1 treated J20 animals (n=8) spent 66% ± 10 of their time in the novel arm (p=0.04). The Y maze data suggests that OXS-N1 resulted in an improvement in hippocampal-dependent short-term memory.

OXS-N1 treated animals had a large, but non-significant, increase in the total number of Dcx+ cells in the dorsal DG (7,894 cells/mm^3^ ± 2,429 (vehicle, n=8) vs. 18,555 cells/mm^3^ ± 4,448 (OXS-N1, n=8; p=0.1 Mann-Whitney test). The increase in Dcx+ cell density was also not significant in the ventral DG (data not shown). A modest non-significant increase was noted in BrdU+Dcx+ cell density in the dorsal dentate gyrus upon OXS-N1 treatment (1,851 cells/mm^3^ ± 680 (vehicle, n=8) vs. 2,488 cells/mm^3^ ± 725 (OXS-N1, n=8; p=0.53) (data not shown). The dentate gyrus was also analysed for the presence of recently generated mature BrdU+NeuN+ neurons. No statistically significant difference between compound treated and vehicle treated animals was found in either dorsal (Figure 5F) or ventral DG (data not shown) when the DG was taken as a whole. However, when the dorsal DG was split into suprapyramidal and infrapyramidal blades, the density of both Dcx+ cells and BrdU+NeuN+ cells in the infrapyramidal blades were significantly greater in the OXS-N1 group compared with the control group (Figure 5G, H). This suggests that there may be an effect of this compound upon the generation of newborn neurons, possibly born during the first two weeks of drug administration, before BrdU was given. This was supported by evidence for an improvement in behavioural scores in the Y-maze as well as a reduction in the locomoter activity seen in J20 mice.

### Transcriptomics analysis after OXS-N1 administration

We commenced mechanism of action and target deconvolution studies by carrying out bulk RNA sequencing on compound treated neurospheres. To mimic our in vitro screen, we exposed proliferating neurospheres to 3 days of OXS-N1 after which we collected cells and prepared RNA (Fig. 6A). In parallel experiments, we exposed cells to OXS-N1for the same period of time but collected cells after 6 days in differentiation media (Fig. 6A).

**Figure 6:**
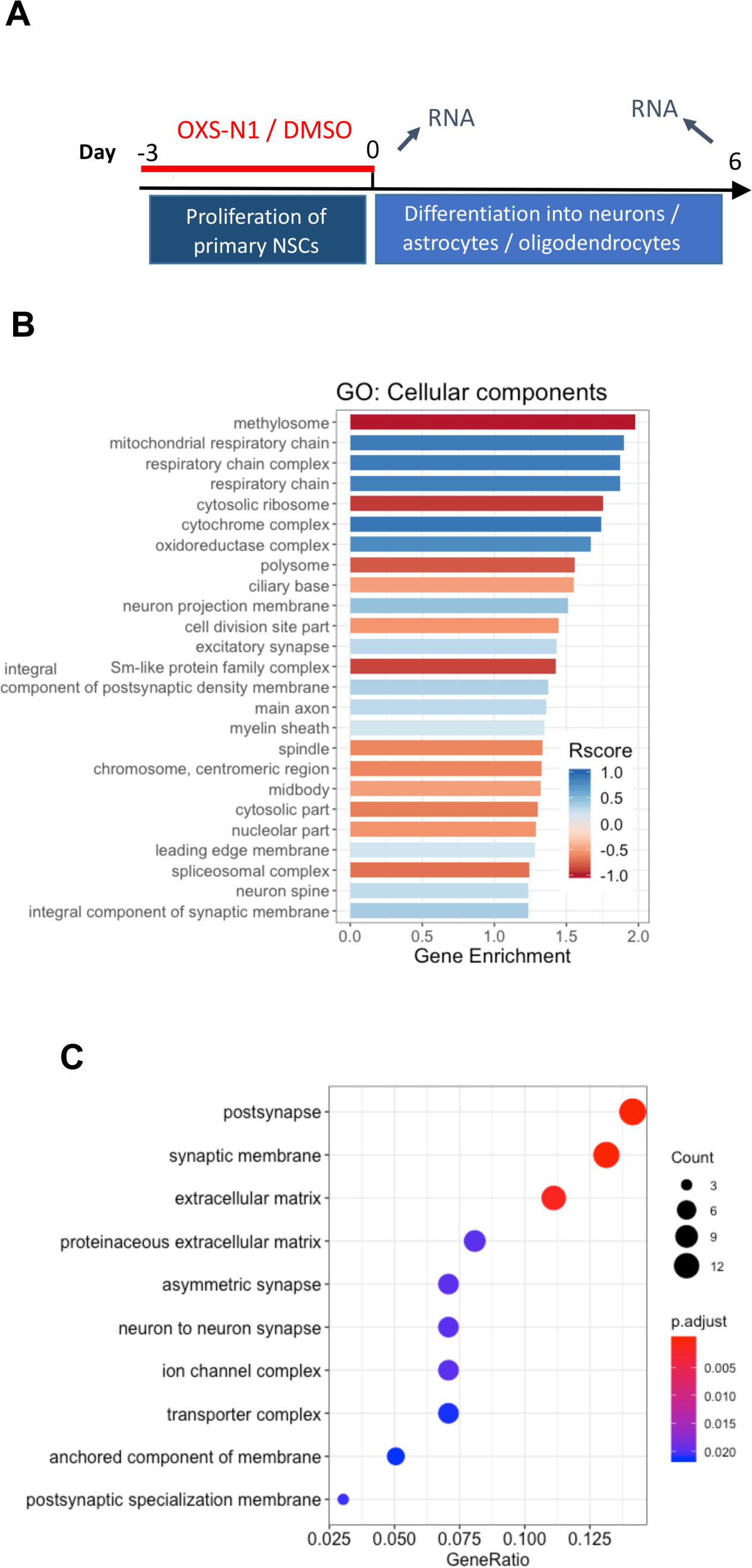
Transcriptomics analysis of neurospheres treated with OXS-N1. (A) Schematic of experimental strategy. (B-C) Gene ontology of The Rscore shiws the proportion of up/down regulated genes in a given pathway. The enrichScore is the ratio of the number of observed over expected genes in a given pathway. Pathways are ranked by their significance, with the most significant at the top.

We first carried out an analysis of treatment, controling for state (proliferation/differentiation) and region (SVZ/SGZ). This approach broadly examined the effect of treatment across all samples, controlling for differences between regions and state. This model incorporates more samples so potentially has greater statistical power. In gene enrichment analyses, we found up-regulation of genes associated with neuronal and cellular respiration-related components (Figure 6B). This is interesting because cellular respiration is known to be upregulated during NSC activation/maturation. Mirroring this, we also found that many down-regulated genes were involved with translational and mitotic machinery, but up-regulation of electron transfer (Figure 6B). In gene ratio analyses, we found associations with the synapse and extracellular matrix and also that the OXS-N1 response was enriched for neurotransmitter molecular functions and the gene activity was localized to brain-associated components (Figure 6C).

Then to begin to understand the effects of OXS-N1 on human cells we caried out a pilot study on hiPSC (Figure S5). After removing the effects of replicates, fraction mapped and the NanoDrop ratio, we found that treatment separated out along PC1, which accounts for 70% of the overall variation (Figure S5A). Many of the genes found to be regulated by OXS- N1 were involved in synapse-related pathways (Figure S5B,C).

## Discussion

### Summary

In the current study we provide evidence for the discovery of a novel pro-neurogenic small molecule. OXS-N1 was identified as pro-neurogenic in a murine DG and SVZ derived neural stem cell (NSC) phenotypic screening assay, in which we tested over 1000 compounds selected from a library based on pharmacophore modelling of known neurogenic compounds. *In vitro* analysis revealed this compound to be capable of increasing neuronal numbers. OXS- N1 was orally bioavailable, stable, penetrated the BBB and was safely tolerated by mice exposed to it for up to 3 weeks. OXS-N1 increased hippocampal neurogenesis *in vivo* after 1 week i.p. or 1 week gavage. It also increased neurogenesis in the SVZ/OB after 3 weeks of gavage. In the J20 model OXS-N1 increased neurogenesis in the infrapyramidal blade of the dentate gyrus after 3 weeks of gavage. We also showed improved performance in the hippocampus dependent Y maze behavioural test in both WT and an AD mouse model after three weeks gavage OXS-N1. We hypothesise that OXS-N1 can improve cognition in animal models of Alzheimer’s disease (AD) via its pro-neurogenic mechanism of action.

### In vitro and in vivo pharmacology

The small molecule OXS-N1 increased neurogenesis in a concentration-dependent manner up to 1µM in both the DG and SVZ NSC-derived differentiation assays. At higher concentrations OXS-N1 still increased neurogenesis compared to vehicle control, but the difference was less marked. Further tests revealed that at higher concentrations OXS-N1 was detrimental to stem cell viability. Thus, the decrease in neurogenic activity of OXS-N1 at higher concentrations could be explained by toxicity-associated effects which is also seen with higher concentrations of the control compound retinoic acid. Pharmacokinetic analysis revealed that OXS-N1 was actively effluxed and had hepatic stability. Upon IP and PO administration in vivo the compound was shown to not only cross the BBB but also to be detectable in the brain and blood for a few hours after administration.

### IP injections of OXS-N1 increased neurogenesis

Animals were initially administered OXS-N1 IP as this was predicted to be the most efficacious route of delivery. Animals were injected with the compound for 1 week followed by a 2-week washout. This timing was used to examine the effects of acute administration while enabling sufficient time for newborn neurons to mature and functionally integrate into the synaptic circuitry and be detectable with immunohistochemistry [51]. Indeed, we found that numbers of newly generated young neurons (BrdU+Dcx+) were significantly increased. Confirming this, we also showed that newly generated neurons exhibiting a mature marker (BrdU+NeuN+) were significantly increased.

When examining the DG, we analysed the dorsal and ventral regions separately. The hippocampus, depending on the dorso-ventral location, has different afferents and efferents which are purported to have different functions [42, 39, 43, 40, 41, 44, 45]. The dorsal hippocampus is is important in memory and spatial awareness whereas the ventral hippocampus has more limbic system inputs, is important in anxiety and stress responses [43–45]. Further strengthening these results, we showed that OXS-N1 effects occurred in the dorsal as well as in the ventral dentate gyrus. These results suggested that OXS-N1 might also affect multiple different aspects of hippocampal-based memory, cognition and emotion in humans.

We demonstrated that one week of IP OXS-N1 administration measurably increased neurogenesis in the hippocampal stem cell niche, however the SVZ appear unaffected as evidenced by proliferation markers (pHH3 and Ki67). The same proliferation markers in the same animals were also not statistically significantly different in the hippocampal stem cell niche even though neurogenesis *per se* was increased. This could be because the effect of OXS-N1 was not to increase proliferation but to increase newborn neuronal survival. It is estimated that 30-70% of DG neuroblasts and circa 50% of SVZ derived neuroblasts die under homeostatic conditions [52, 53]. Future studies may show that in the SVZ OB neurogenesis is in fact increased in these mice when measured with BrdU, Dcx and NeuN. As will be discussed below, this is what we found in WT mice after the three weeks of OXS-N1 gavage.

### Oral administration of OXS-N1 increased neurogenesis and increased memory

With 1-week oral OXS-N1 administration, we demonstrated significantly increased hippocampal neurogenesis. Similar to the 1-week IP group, increased neurogenesis after OXS-N1 was seen in the dorsal and ventral hippocampus, and with BrdU+Dcx+ as well as BrdU+NeuN+ counts. The combination of these two results demonstrates that short regimens of OXS-N1 increase neurogenesis. However, we did not see effects of 1 week OXS-N1 gavage in a hippocampal dependant Y maze. This may be explained by the newborn neurons not having a long enough period for effective maturation and proper synaptical integration to contribute to hippocampal functionality.

However, with the longer 3-week gavage dosing paradigm we did see an effect at the behavioural level, with evidence from the Y-maze that OXS-N1 increases memory. It takes around 3 weeks for newborn hippocampal neurons to integrate [54–56]. This timing would thus give newborn neurons enough time to integrate functionally into the hippocampal circuitry and elicit measureable behavioural changes. To accommodate the memory and learning requirements of the individual, newborn neurons either integrate or apoptose as part of normal regulation of the niche.

Surprisingly, the 3-week gavage paradigm did not show increased hippocampal neurogenesis, suggesting that OXS-N1 may only have transient neurogenic effects. However, there was evidence for increased OB neurogenesis after three weeks of OXS-N1. More young newborn neurons were found in the core of the OB and more mature newborn neurons were found in the granular layer. Newborn neurons in the olfactory bulb are generated in the SVZ and migrate along the rostral migratory stream to their target destination. Neuroblasts originating from the SVZ have been shown to have a peak of BrdU+ cell density/number in the OB 1 month after BrdU injection [57, 52]. The longer (3-week) administration regime may have been able to demonstrate additional new-born neurons in the OB precisely because there was more time during the drug administration as well as during the longer wash-out period for neurons to migrate into and settle in the olfactory bulb. The fact that SVZ neurogenesis was increased in this paradigm adds strength to our contention that OXS-N1 is pro-neurogenic.

### OXS-N1 effects on neurogenesis in an Alzheimer’s disease model

Adult human hippocampal neurogenesis has recently been confirmed with immunohistochemistry and single cell RNA sequencing approaches [58, 16–18]. Interestingly Alzheimer’s disease has been associated with decreased hippocampal neurogenesis [58, 59]. It has been suggested that hippocampal neurogenesis is impaired in J20s at 4-11 weeks but not at 14 weeks [60]. Reductions in immature neurons in the SVZ and SGZ of J20s has also been observed at 3 months of age up until 12 months [29]. This matches our finding that Dcx+ immature neurons were very rare in the J20 brains we analysed, as were cells that took up BrdU during the one-week exposure, therefore we only quantified Dcx+ cells. There is also consistent evidence that J20s have impairments in hippocampal specific tests such as the Morris water maze (MWM), Y-maze and novelty object recognition (NOR) [61, 48, 50].

They also exhibit a propensity to hyperactivity in the open field test (OFT) and elevated plus maze [48–50]. Our finding that OXS-N1 decreased locomotion and increased short-term memory in the Y maze indicated that it has an effect in this model of Alzheimer’s disease.

Increased locomotion can in and of itself increase hippocampal neurogenesis, however the compound decreased locomotion, suggesting that the increased neurogenesis was not due to locomotion. We found strong trends towards overall increased neurogenesis which would likely be statisticaly significant with a larger cohort. The large variation in response is likely explained by the fact that individual J20 mice of a given age are variably affected.

Furthermore, because of the strong trend we queried if subregions of the dentate gyrus had different OXS-N1-induced effects. The infrapyramidal blade in the dorsal hippocampus did show statistically significant increased neurogenesis as measured by both Dcx+ and BrdU+NeuN+ cell numbers. There are several phases of AD and large variety of symptoms elicited in animal models of the disease. Thus, it will be important to determine how broadly OXS-N1 may increase neurogenesis.

Neurogenesis has also been extensively investigated in animal models of AD. In general, single human mutations associated with AD are associated with reduced neurogenesis in mouse models. For example, Tg2576, PrP–APPSwe and PDGF–APPInd all have reductions in proliferation and survival of SGZ cells [62, 63]. A variety of studies have been performed in PDGF-APPSwe,Ind (also known as J20) mice, with conflicting results. Amyloid accumulation and plaque formation in J20s begins from as early as 2 months and by 7 months almost all mice show evidence of amyloid plaques [64], although others report that amyloid plaques only become detectable reliably after 5 or even 9 months of age [49]. Neuronal loss in the hippocampal CA1 field, or reduction in dentate gyrus volume – both hallmarks of the human disease – have been reported from as early as 12 weeks of age [48, 49]. Neurogenesis has been assessed in J20s in several studies, using slightly different methodologies. The number of proliferative Ki67+ cells in the DG decreases with age, both in wild-type and J20 mice, but whether J20s have reduced neurogenesis compared to wild-type controls may depend on age. Fu and colleagues report fewer Ki67+ cells in DG of young (4-6 week old) J20s compared to controls but more in aged animals (18 month old) [60]. Others report increased DG neuronal precursor proliferation as assessed by BrdU incorporation in J20s compared to wild-type controls at both 3 months and one year of age [47, 61]. Conversely, while Lopez-Toledo and colleagues also report more BrdU+ cells in DG of 3-month-old J20s compared to controls, they find a decrease in BrdU+ cells in J20s at all later stages studied [65].

### How does OXS-N1 work?

Adult neurogenesis comprises a large range of cellular events from stem cell quiescence/activation to synaptic integration, and these phases are governed by waves of gene regulatory networks. Our data indicated that OXS-N1 did not activate stem cells as evidenced by BrdU+GFAP+ cells. We also had evidence that it does not increase proliferation of intermediate progenitor cells as measured by Ki67+pH33+. Thus, it may be that OXS-N1 affects the later stages of neurogenesis. Our bulk transcriptomics analyses of neurospheres exposed to OXS-N1 revealed several interesting clues as to how may work. Firstly, there were several cellular components regulated by OXS-N1 indicative of metabolism such as “mitochondrial respiratory chain”, “respiratory chain complex”, “cytosolic ribosome”, etc. This is interesting since tight regulation of metabolism is essential for lineage progression in adult brain stem cells [66]. The other major set of clues that emerged from the neurosphere analysis were terms associated with the synapse, such as “postsynapse”, “synaptic membrane”, “asymmetric synapse”, etc. This was reproduced by the gene ontology terms that emerged from our preliminary analysis of human iPSC cells which were exemplified by “postsynapse”, “postsynaptic membrane”, “synaptic membrane”, “dendritic spine” etc. Increasing synaptic contacts of newborn neurons would help them integrate into adult circuitry and therefore likely improve both survival and function. We have also found evidence in parallel studies for OXS-N1 working through the protein vimentin. Future work will help reveal the exact mechanism(s) of action of OXS-N1 in increasing neurogenesis.

### Conclusion

We show here that a primary cell culture-based phenotypic screen of a small molecule library identified a compound that could increase neurogenesis and improve performance in behavioural studies in mice. Given the aging of the human population, the preponderance of AD and the lack of effective medications for it, there is a dire need for novel types of therapies. This study supports the concept that increasing numbers of newborn neurons should continue to be explored as a novel form of therapy.

## SUPPLEMENTARY MATERIAL

### Supplementary Methods

#### In Silico pharmacophore modelling

A neural stem cell modulator compound database was established using the ChemBioFinder program (ChemBioOffice Software package, PerkinElmer Inc.), and the ChemBioDraw program (PerkinElmer) was used to draw structures of compounds as they were entered into the database. Conversion of chemical structures to 3D models and energy optimization for database mining and phamacophore creation was carried out using ChemBio3D (PerkinElmer). Phamacophores were generated using FieldTemplater (Cresset). Phamamcophore and compound-based similarity scans were carried out in Forge (Cresset), protonation at physiological pH was used and the scans were carried out at 100 conformations per compound and results analysed within Forge. Structure- based similarity screens were carried out using Knime (Konstanz information miner, KNIME.com AG) using CDK (Chemical Development Kit, open source) and Pipeline Pilot (Scitegic) and results were analysed using ChemFileBrowser (Hyleos).

A searchable neurogenesis modulator database was created in order to screen compound libraries. Information on commercially available compounds known to interact with pathways that influence NSC differentiation and/or proliferation (Wnt/β-catenin signalling, Sonic Hedgehog signalling, EGFR, FGFR, MAPK) was assimilated into the library. Using this, a library of 351 compounds was generated with all compounds having been shown to have *in vitro* or *in vivo* effects on neurogenesis based on the literature. These compounds were used to populate lists of chemically similar compounds within the target databases using the Tanimoto algorithm-based (structure-based) processing setups in the Knime and Pipeline Pilot programs.

From the selection of neurogenic compounds, groups that had the same target were selected to generate pharmacophores using FieldTemplater. These were chosen as initial testing compounds to be used in the NSC screens. A further 1000 compounds were also selected to be prioritized in *in vitro* assays. The remaining compounds were selected randomly from our library of compounds to increase the richness of the chemical matter. SDF Reader nodes contained the screening library and query compound for the top and bottom nodes, respectively. A new query compound was set for every screen. The Fingerprints nodes converted the chemical structures in the SDF files into Daylight-type fingerprints (series of vector sets). The Fingerprint Similarity node compared the query compound’s fingerprint to the fingerprint of every compound in the library. Outputs for sorting and analysis were written in SDF and Excel table formats by the SDF Writer and XLS Writer nodes, respectively.

---

**Figure S1.**
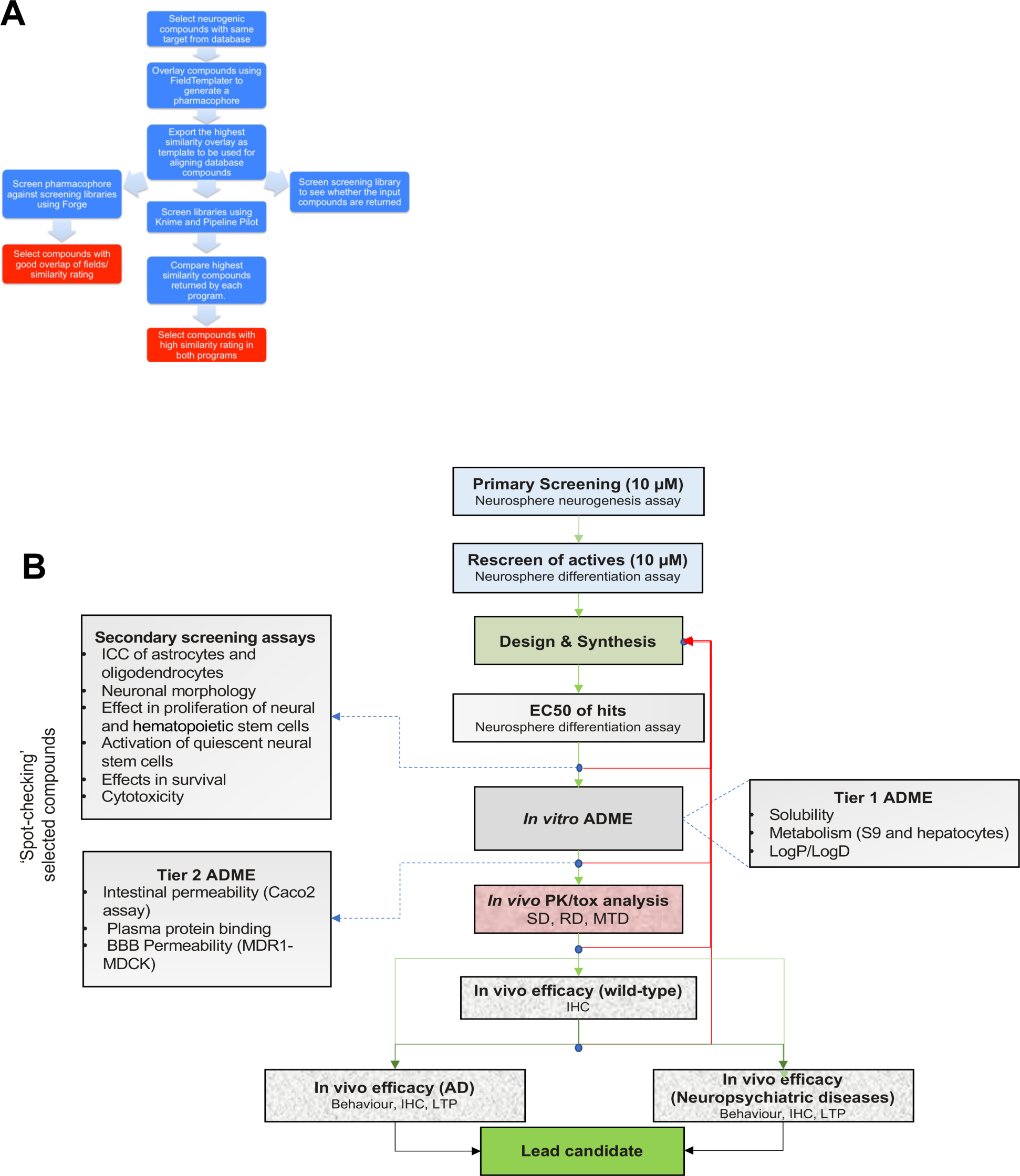
Screening Strategies. A: Sceening and selection of compounds for in vitro testing. B: Screening cascade for in vitro, structure activity relationships and in vivo screeing to lead candidate.

**Figure S2.**
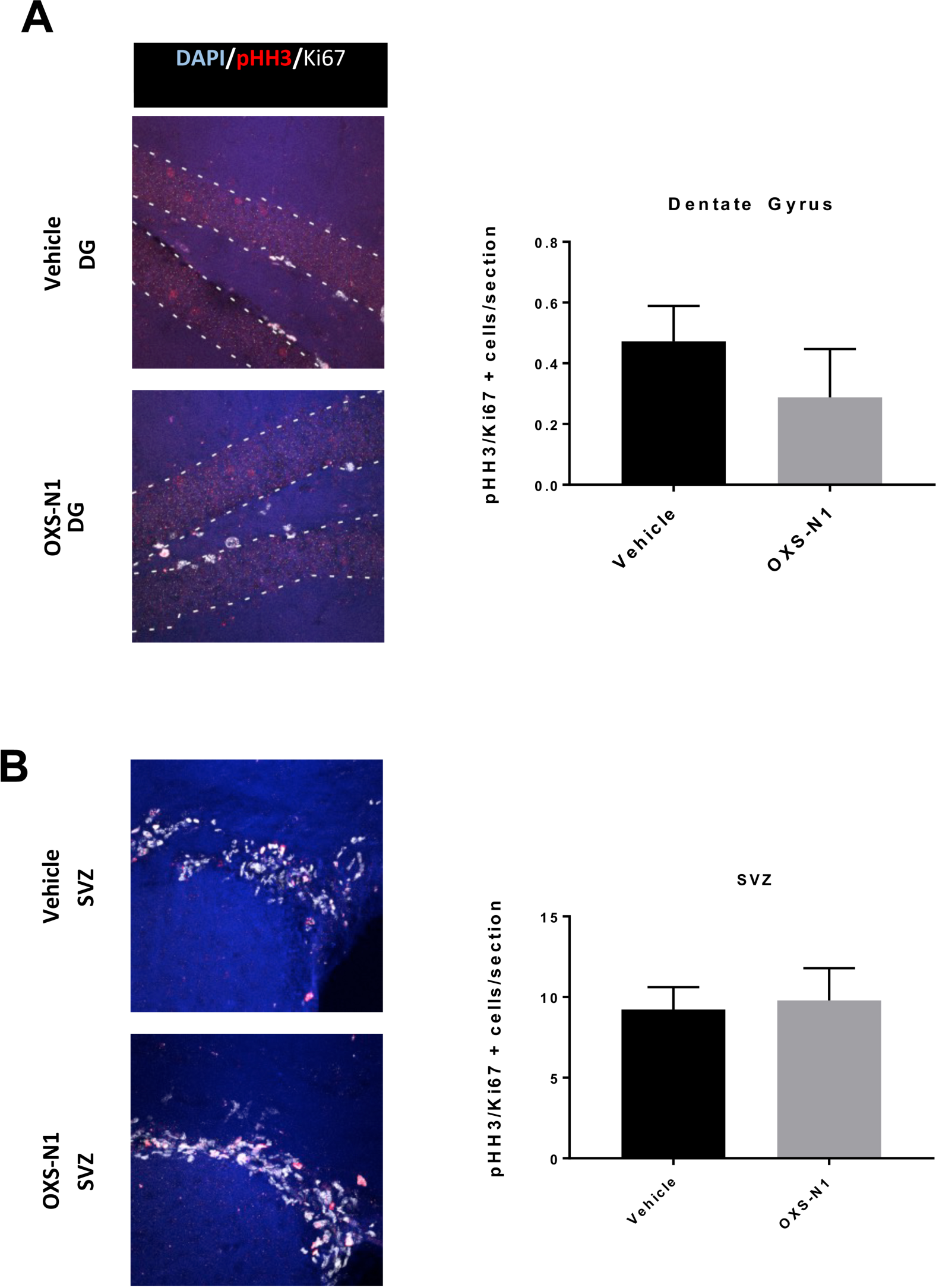
Proliferation in the DG and SVZ after OXS-N1. A: pHH3 (red) and Ki67 (white) cells in the DG after 1 week IP OXS-N1 administration. B: Analysis of pHH3+Ki67+ cells in the SVZ of animals treated with OXS-N1 for 1 week IP.

**Figure S3.**
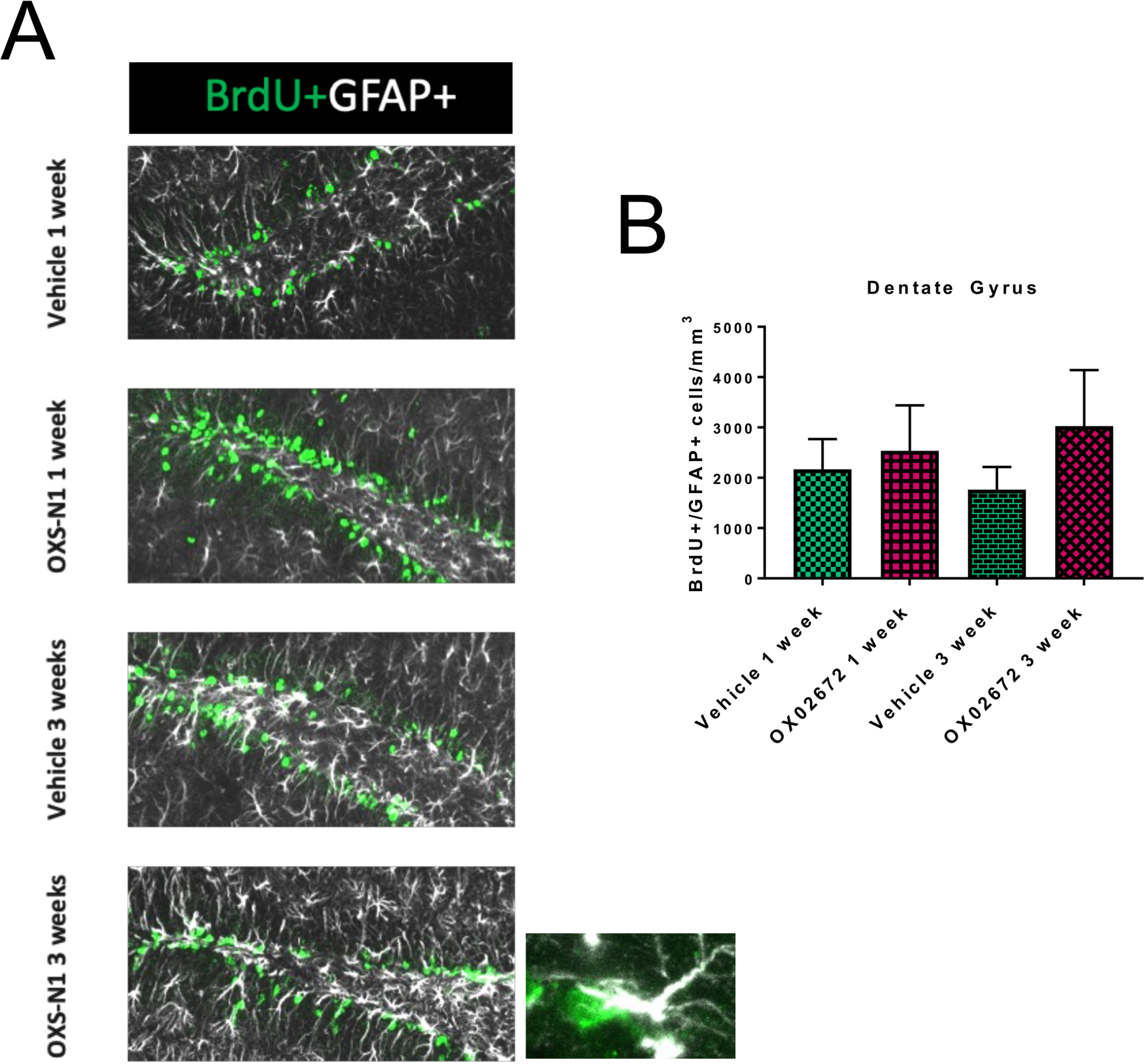
Dentate gyrus stem cells after OXS-N1. A: BrdU (green) and GFAP (white) positive cells in the DG of animals treated with OXS-N1 orally for 1 or 3 weeks. B: Analysis of Brd+GFAP+ cells in the DG of animals treated for 1 or 3 weeks with

**Figure S4.**
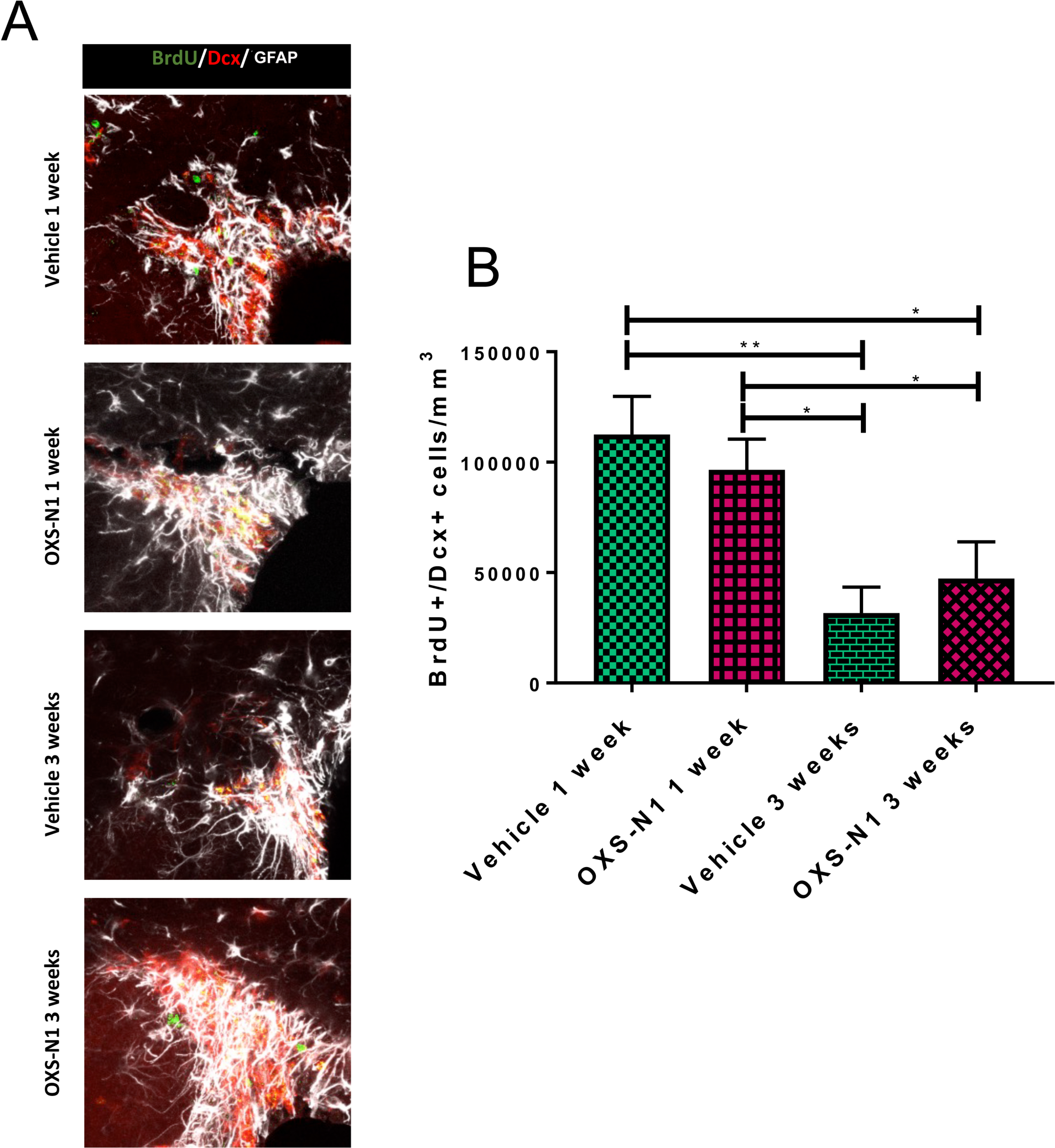
Newborn neurons after oral OXSN-1. A: SVZ of animals treated with OXS-N1 orally for 1 or 3 weeks showing BrdU (green), Dex (red) or GFAP(white) immunohistochemistry. B: Analysis of BrdU+Dcx+ cells in the SVZ of animals treated with OXS-N1 by gavage for 1 and 3 weeks. Note the significant differences bewteen the 1 week and 3 week groups but lack of effect due to OXS-N1 administration. One way ANOVA, *p<0.05, **p<0.01

**Figure S5.**
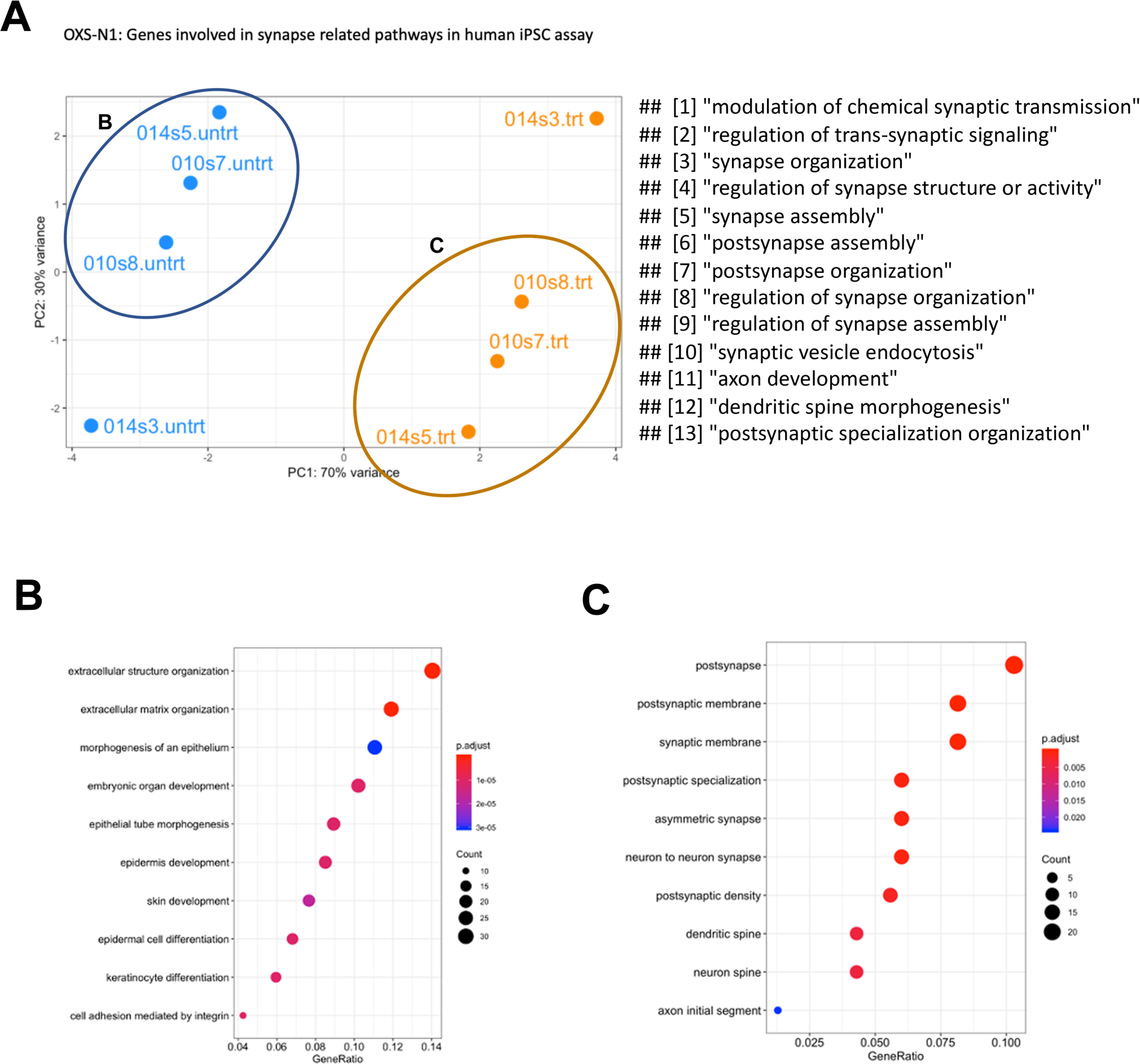
Transcriptomics carried out on hlPSC from 2 healthy human cases. A: principle componnt analsys shows that that treatment accounted for most variation. Samples encircled are shown in more detail in **B** and C.

